# Spontaneous activity competes externally evoked responses in sensory cortex

**DOI:** 10.1101/2020.08.18.256206

**Authors:** Golan. Karvat, Mansour Alyahyay, Ilka Diester

**Affiliations:** Optophysiology Lab, Institute of Biology III, University of Freiburg, Freiburg, 79104, Germany; Bernstein Center Freiburg, University of Freiburg, Freiburg, 79104, Germany; BrainLinks-BrainTools, University of Freiburg, Freiburg, 79104, Germany

**Keywords:** LFP, oscillation, beta-burst, resting-state-network, default-network, detection, real-time, cortex, forepaw, somatosensory

## Abstract

The functional role of spontaneous brain activity, especially in relation to external events, is a longstanding key question in neuroscience. Intrinsic and externally-evoked activities were suggested to be anticorrelated, yet inferring an antagonistic mechanism between them remains a challenge. Here, we used beta-band (15-30 Hz) power as a proxy of spontaneous activity in the rat somatosensory cortex during a detection task. Beta-power anticorrelated with sensory-evoked-responses, and high rates of spontaneously occurring beta-bursts predicted reduced detection. By applying a burst-rate detection algorithm in real-time and trial-by-trial stimulus-intensity adjustment, this influence could be counterbalanced. Mechanistically, bursts in all bands indicated transient synchronization of cell assemblies, but only beta-bursts were followed by a reduction in firing-rate. Our findings reveal that spontaneous beta-bursts reflect a dynamic state that competes with external stimuli.

## Introduction

The brain is constantly active, even at resting-states in the absence of external stimuli (Deco et al., 2013). Spontaneously active resting-state networks (RSN) were found in memory, visual, auditory, tactile and sensorimotor regions, with activity patterns similar to task-evoked responses (Afrashteh et al., 2020; Damoiseaux et al., 2006). Functional connectivity studies in humans suggest that a default network, spontaneously activated at resting-state and deactivated upon increased cognitive demands, antagonizes a network involved in active attention to external sensory input (Biswal et al., 1995; Fox et al., 2005; Fransson, 2005; Raichle et al., 2001; Raichle, 2015). However, weather these networks anticorrelate each other is under debate, and their antagonizing mechanisms and influence on local circuits remain unknown (Buckner and DiNicola, 2019).

Here, we utilized the rate of high-power bursts in the beta-band (15-30 Hz) of local-field-potentials (LFP) as an indicator of spontaneous activity to investigate its influence on detection in real-time. Several lines of evidence relate the RSN to beta-bursts. First, spontaneous correlated oscillatory activity in beta (termed ‘beta-connectome’ (Seedat et al., 2020)) was reported in anatomical regions corresponding to RSN (Brookes et al., 2011; Hipp et al., 2012). A recent study derived this beta connectome from burst rate (Seedat et al., 2020). Second, beta oscillations are dominant during resting state (Panagiotaropoulos et al., 2013) and bursts are responsible for virtually all beta-band power modulation (Feingold et al., 2015). Third, task-dependent desynchronization of beta was observed in the somatosensory (Bauer et al., 2006; van Ede et al., 2010), visual (Chen et al., 2017), auditory (Pesonen et al., 2006), and motor (Kilavik et al., 2013) cortices, resembling RSN deactivation (Biswal et al., 1995; Raichle et al., 2001). The task-dependent averaged power-modulation was attributed to changes in burst-rates in rodents, non-human-primates and humans (Feingold et al., 2015; Shin et al., 2017; Tinkhauser et al., 2020). Fourth, the burst duration (50 to a few hundred ms (Karvat et al., 2020)) is similar to ‘packets’ of neural activity, which are conceived as messages initiated in a particular cortical region and spread as a wave over the cortex. Most of these packets are generated spontaneously, and spontaneous and sensory-evoked packets are remarkably similar (Luczak et al., 2015).

We found that bursts in all bands indicate transient synchronization of flexible neuronal-networks, but only beta-bursts were followed by a reduction in population firing-rate. High rates of beta-bursts predicted reduced detection, and this effect can be counterbalanced bi-directionally in real-time by adjusting the stimulus intensity according to burst-rate.

## Results

### Rats detect forepaw vibration

To investigate the relationship between externally and internally generated activities in detection, we developed a novel detection task in rats (Fig. 1A, Fig. S1 and methods). To allow future translation of our findings to writing-in sensations into the brain needed for neuro-prosthetics, we targeted the forepaws of the rats with a vibro-tactile stimulus. The freely moving rats pulled and held a lever and released it upon detection of vibration generated by a piezo actuator. As unimanual and bimanual actions are represented differently in cortex (Donchin et al., 2001), we ensured unimanual manipulation of the lever by enclosing it between two walls. The rats could choose which paw to use, and 37.5% (3 out of 8 rats) preferred the left paw. Since beta-activity is prominent during periods of waiting for stimuli (Kilavik et al., 2013), we introduced a holding duration prior to the stimulus. To prevent the rats from using a timing strategy, this holding-period was pseudo-randomly defined. Trained rats were able to hold the lever steadily for 600, 1600 or 2600 ms (Fig. 1B-C and Fig. S1), and to respond to the 300 ms, 300 Hz, 28 μm sinusoidal lever vibration within 369.7 ± 6.3 ms (mean ± 95% confidence interval, N=40 sessions, Fig. 1D and Fig. S1). To prevent the use of the piezo sound as cue, in 20% of trials we vibrated a control lever located in the cage but out of the rats’ reach. In these control trials, the rats had to keep holding the lever >620ms to obtain a reward. The response rate of trained rats to vibration was significantly higher than to the control (Fig. 1E, Wilcoxon *z*=7.66, *p*=1.93*10^−14^), yielding a d’ value of 1.56 ± 0.25 (mean over sessions ± 95% confidence interval).

**Fig. 1.**
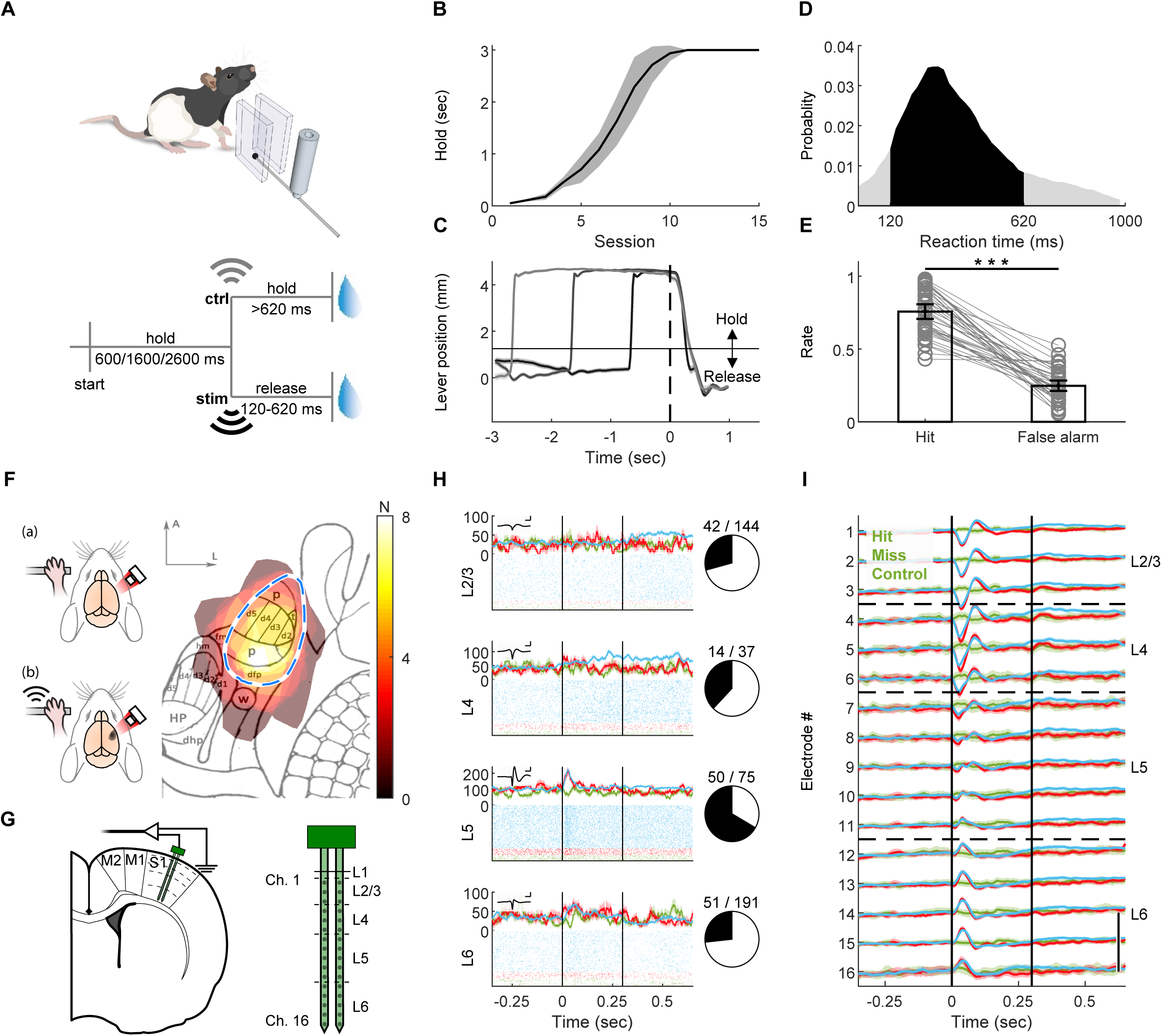
Rats detect forepaw vibration. (A) Upper panel: simplified illustration of the behavioral setup. A lever was enclosed between two walls and could be moved on the horizontal dimension. Piezo actuator clamped to the lever conveyed the vibrotactile stimulus. For more details, see Extended Data Fig. 1. Lower panel: task design. To start a trial, the freely moving rats had to pull the lever in their preferred direction and hold it for one of three pseudorandom durations. Upon vibration (stim), the rats had to release the lever within 120-620 ms to gain a sucrose water reward. In 20% of the trials, an actuator located in the chamber but clamped to a lever out of the rat’s reach was activated (ctrl). The rat had to keep holding for longer than 620 ms in order to gain a reward. (B) Holding training progress. Mean ± 95% confidence interval of the holding duration of 8 rats. Rats had to hold the lever passed the threshold position for at least the holding time to obtain a reward. (C) Rats learned to hold the lever steadily and respond to the cue. Mean ± 95% confidence interval of the lever position (measured by the angle encoder) at “hit” trials. Time *t*=0 indicates the stimulus onset. Horizontal line at ∼1mm indicates the hold/ release threshold. Trials with holding time = 600 ms are indicated in black, 1600 ms in gray and 2600 ms in light grey. (D) Histogram of reaction (release) times in stim trials. Vibration was presented from *t* = 0 to *t* = 300 ms. Black area: RTs within the allowed response window. Gray: RTs of early or late release trials. N=40 sessions from 8 rats. (E) Hit rate – ratio of trials with lever release within <620 ms after real stimulus onset, from all vibration trials. False alarm rate – ratio of responded trials within <620 ms after control actuator onset from all control trials. Bars indicate the mean and error bars the 95% confidence interval. Gray circles and lines – individual sessions. ***-p<0.001, Wilcoxon rank sum test. (F) Intrinsic signal imaging. Left panel: Schematic illustration of procedure. (a) The brains of lightly anesthetized rats were imaged through the thinned skull under 855 nm illumination. Upon vibrotactile stimulation of the forepaw (b), responsive cortex absorbed more light compared to blank trials. See methods for details. Right panel: responsive areas of 8 rats overlaid on projected published somatotopic map^26^. N (color bar) indicates number of rats. Blue dashed line: assumed SI forepaw location based on map. p, palm; t, thumb (pollux); d1-5, digits; dfp, dorsal forepaw; fm, forelimb muscle; hm, hindlimb muscle; w, wrist whiskers; dhp, dorsal hindpaw; HP, hindpaw; Scalebar: 1 mm. A, anterior; L, lateral. (G) Left panel: schematic illustration of implantation locations of the 2-shank, 32-channel probe. S1: primary forelimb somatosensory cortex. M1: primary motor. M2: secondary motor. Dashed lines: estimated borders between cortical layers. Right panel: magnified illustration of the probe, indicating the location of channel relative to surface and tip and estimated borders between cortical layers as shown in **H** and **I**. (H) Left panels: Peri-stimulus-time-histogram (PSTH, top) and raster plots (bottom) of representative single units in Layer 2/3, 4, 5 and 6. Blue: hit. Red: miss. Green: control. Inset: averaged waveform of the unit. Scalebar: 1 ms / 250 μV. Right panels: the ratio of modulated units (black) from total single-units. (I) Mean ± 95% confidence interval of the LFP color coded as in **H**. Vertical black lines indicate stimulus onset and offset (*t*=0 to 0.3 sec). Scale bar: 200 μV. See also Fig. S1 and S2.

### Evoked activity encodes the presence, not the detection, of a sensory cue

We investigated the cortical response to vibration via chronically implanted multi-electrode laminar probes. To ensure accurate positioning while avoiding extensive bleeding from surface vasculature, we developed a survival intrinsic-signal-imaging (based on cerebral blood-flow (Grinvald et al., 2016)) and implantation surgical procedure for rats (Fig. 1F-G, Fig. S2 and methods). With this approach, we were able to take differences in vasculature and somatotopy between individuals into account which allowed a more precise positioning of the probes compared to estimations based on classic somatotopic maps (Chapin and Lin, 1984) of primary somatosensory cortex (SI) location.

We further validated the insertion location by detecting stimulus evoked modulation of single-units firing-rate (Fig. 1H) and LFP (Fig. 1I). The LFP showed depolarization followed by hyperpolarization in dorsal electrodes and hyperpolarization in ventral electrodes. Analysis of the event-related-potentials (ERP, Fig. 2A) and current-source-density (CSD, Fig. S2B) immediately following the stimulus revealed the strongest response at electrodes located 600-800 μm below the surface. This response peaked 25 ms after stimulus onset, as expected from layer 4 (L4) of SI receiving a direct thalamo-cortical input (de Lafuente and Romo, 2006; Narayanan et al., 2017). Furthermore, histology (Fig. S2C) confirmed the position of these specific electrodes at the granular layer.

**Fig. 2.**
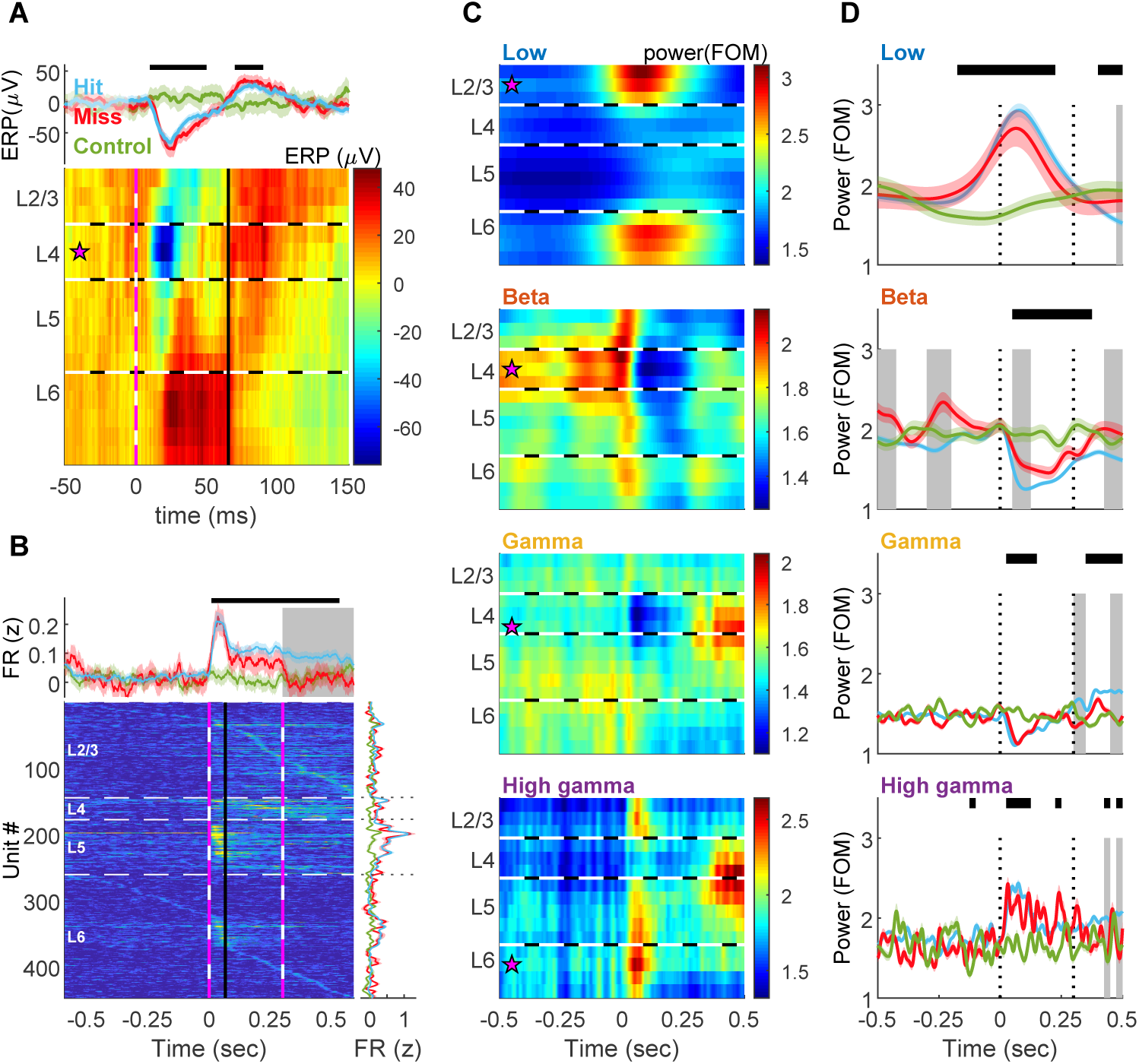
Band and layer differences in detection. (A) Event related potential (ERP), calculated as the mean of raw LFP aligned to stimulus onset (time *t*=0). Upper panel: mean ± 95% confidence interval on electrode 5 (located in L4) averaged over hit (blue), miss (red) and control (green) trials. Top black bar indicates significant (*p*<0.01) difference between hit and control trials, in ANOVA for repeated measures with Tukey’s critical value for multiple comparisons. Main effect of time: *F*_9,11646_= 34.6, *p*= 6.95*10^−61^, time × behavior: *F*_18,11646_= 11.93, *p*= 2.63*10^−35^. Lower panel: the ERP on each electrode at hit trials. Magenta star: the electrode used in the upper panel. Horizontal black and white dashed lines indicate the estimated borders between layers. Vertical pink and white dashed line designates stimulus onset (*t* = 0). Vertical black line denotes phase transition from de-to hyper-polarization in L4 at hit trials (blue trace in upper panel passes 0 after the trough, *t* = 0.065).N=10 sessions from 5 rats. (B) Upper panel: mean ± 95% confidence interval of the z-normalized firing-rate (FR) of all units (N=447 units from 5 rats), color coded as in **A**. Piezo activation started at *t*=0 and lasted 0.3 sec (green dashed lines). Top black bar indicates significant (*p*<0.01) difference between hit and control trials, in ANOVA for repeated measures with Tukey’s critical value for multiple comparisons. Gray background indicates a significant (*p*<0.01) difference between hit and miss. Main effect of time: *F*_40, 53520_= 25.6, p= 1.14*10^−186^, time × behavior: *F*_80,53520_= 9.47 *p*= 2.03*10^−82.^ Lower left panel: FR during hit trials. Units are sorted according to layer and timing of maximal FR. Horizontal black and white dashed lines indicate the estimated borders between layers. Vertical pink and white dashed lines designates stimulus onset (*t* = 0) and offset (*t* = 0.3). Lower right panel: mean FR ± 95% confidence interval of each unit at time *t*=0 to *t*=0.065 sec (depolarization phase of the ERP at L4, denoted by black line on left). Color coded as in **A**. For all layers except L2/3 the FR was significantly higher at hit trials compared to control (*p=* 0.28, 5.8*10^−3^, 2.24*10^−7^ and 0.038 for layers 2/3, 4, 5 and 6, respectively, 2-way ANOVA with Tukey’s post-hoc). There were no differences between hit and miss trials (*p*= 1.0 in all layers). The FR in L5 was significantly (*p*< 3.49*10^−4^) higher than in all other layers. (C) Induced power spectrograms of hit trials, computed by transforming the LFP to the frequency domain using Morlet wavelets, normalizing to the median per frequency and averaging over hit trials. Low: 3-10 Hz. Beta: 15-30 Hz. Gamma: 40-90 Hz. High gamma: 95-120 Hz. Magenta star: most modulated channel used in **D**. FOM-fraction of the median. (D) Comparison of the power dynamics of each band in the different behavioral categories. Behavioral color coding (hit, miss and control), black bars and gray background are the same as in **B**. Main effect for intersection time × behavior: low: *F*_78,43407_= 13.13, *p*= 1.24*10^−162^; beta: *F*_78,43407_= 2.66, *p*= 1.24*10^−13^; Gamma: *F*_78,43407_= 3.43, *p*= 1.57*10^−22^; High gamma: *F*_78,43407_= 1.89, *p*= 3.46*10^−6^.Differences between hit and miss trials prior to and during stimulus (0-0.3 sec, dotted black lines) were observed only in beta. See also Fig. S2-S5.

The ERP was stimulus dependent, as it was absent in control trials. However, it was detection independent, as no differences were observed between hit and miss trials (Fig. 2A, upper panel). The population firing rate (FR) exhibited similar stimulus-evoked dynamics. Upon stimulus presentation, but not activation of the control piezo, the population FR exhibited a robust increase (Fig. 2B and Fig. S3A-B). This response was observed in units across all layers and was strongest in layer 5 (L5). The population FR in SI was shown to drive higher cortical areas to detect a stimulus (Romo and Rossi-Pool, 2020). Nonetheless, during the depolarization phase of the ERP (0-65 ms, Fig. 2A) the stimulus-induced response in FR was similar in hit and miss trials (Fig. S3C). The FR differed between hit and miss after 65 ms in L2/3 and after stimulus offset (300 ms) in L4, plausibly due to the movement towards the reward spout, since forepaw motor and sensory cortices overlap in rodents (Sanderson et al., 1984). Hence, the existence of a stimulus, but not its detection, is encoded by FR and ERP in SI. This raises the question, whether other neuronal signals are associated with detection in SI.

### Band and layer differences in detection

LFP-oscillations are a putative neuronal signal to be associated with detection in SI. Neuronal networks oscillate in a broad range of frequencies, and these oscillations are an effective mechanism to generate temporal synchrony (Buzsáki and Draguhn, 2004). Neural oscillations are divided into frequency-bands (here termed low [3-10 Hz], beta [15-30 Hz], gamma [45-90 Hz] and high-gamma [95-120 Hz]), and categories according to the response to an external stimulus (David et al., 2006): evoked responses are time- and phase-locked to the stimulus, and computed as the power of the ERP in the frequency domain (Fig. S4A). Induced responses are time-locked, but not phase-locked to the stimulus, and extracted as the power averaged over trials (Fig. 2C and Fig. S4B).

We found a relationship between bands, cortical depth, and the stimulus-related categories. Induced power in low frequencies was highest in superficial and deep layers (Fig. 2C), and strongly correlated with the amplitude of the evoked response (power in 3-10 Hz vs. power of the ERP, Pearson’s ρ=0.97, *p*=0, Fig. S5D). The activity in high-gamma was strongest in supra- and infra-granular layers at stimulus onset and L4/5 during movement (Fig. 2C). The high-gamma power strongly correlated with the population firing rate (ρ=0.86, *p*=1.07×10^−287^ in supra-granular, Fig. S5D, and ρ=0.82, *p*=1.33×10^−238^ in infragranular), indicating a close relationship between FR and the induced oscillatory response in the high gamma range. Both the low-frequency evoked response and the high-frequency induced response did not differ between hit and miss trials prior to and during the stimulus (Fig. 2D).

A third and less understood LFP oscillation category, refers to internally driven oscillations (Canolty et al., 2012; David et al., 2006). These ongoing oscillations are thought to represent spontaneous neuronal activity (Hipp et al., 2012). In the gamma-band, spontaneous activity in L4 decreased upon stimulus presentation (Fig. 2C), yet with no difference between detection outcomes (Fig. 2d). Like high-gamma (95-120 Hz), gamma activity in the range of 45-90 Hz differed between hit and miss trials during movement. In the beta-band, the intrinsically-generated activity was strongest in L4. Upon stimulus presentation beta-power decreased. The beta-power was significantly higher in miss trials before and during stimulation (Fig. 2D). Consequently, beta-power was anti-correlated to the FR (ρ= −0.66, *p*=8.57×10^−12^, Fig. S5D).

### Real-time impact of beta on detection

Collectively, the data presented in Fig. 2 indicate that while the presence of the sensory input is associated with modulations of FR, ERP and power in low and high frequencies, detection is associated with the state of intrinsically-generated oscillations in the beta-band. However, the power analysis so far was based on averaging over trials. In order to better understand the relationship between brain signals and behavior, it is important to reveal their trial-by-trial association (Stokes and Spaak, 2016). Since the frequency of occurrence and the duration of brief transient synchronization (bursts) correlate strongly with power modulation (Feingold et al., 2015) (Fig. S5), we used burst-rate as a proxy for the power of ongoing beta activity in individual trials (Shin et al., 2017; Tinkhauser et al., 2020). We categorized trials according to their pre-stimulus burst-rate and found that in trials with high burst-rate the success-rate was significantly lower (Fig. S6A) and the reaction-time (RT, Fig. S6B) significantly higher.

To further establish the impact of beta-bursts on perception, we sought to use their occurrence as the independent variable and assess the dependent behavioral outcome. Since the bursts are highly heterogeneous (Karvat et al., 2020), direct brain stimulation is not expected to mimic the naturally occurring signal. To overcome this obstacle, we developed a system for real-time burst-detection (Karvat et al., 2020). It enabled us to estimate the rate of spontaneously occurring beta-bursts in each trial and to control the timing and amplitude of the vibrotactile stimulus (Fig. 3A). To facilitate amplitude modulation with the best resolution possible in our system (2 μm), we set the standard value per rat as the amplitude resulting in d’≈1 at trials devoid of bursts during the holding time (8.8 ± 1.36 μm, d’=0.99 ± 0.18, mean ± 95% confidence interval, n=5 rats). Confirming our offline findings (Fig. S6), in trials with high burst-rate, rats had lower success-rate and longer RT (Fig. 3C-D). Importantly, in line with our hypothesis of the impact of beta-bursts on detection, increasing the vibration amplitude by 2 μm rescued performance in high-burst-rate trials. This adjustment effect was bi-directional, as in trials with no bursts in the holding-time decreasing the vibration amplitude by 2 μm resulted in detection performance similar to high-burst-rate trials. Taken together, these results demonstrate a direct impact of pre-stimulus beta burst-rate on detection.

**Fig. 3.**
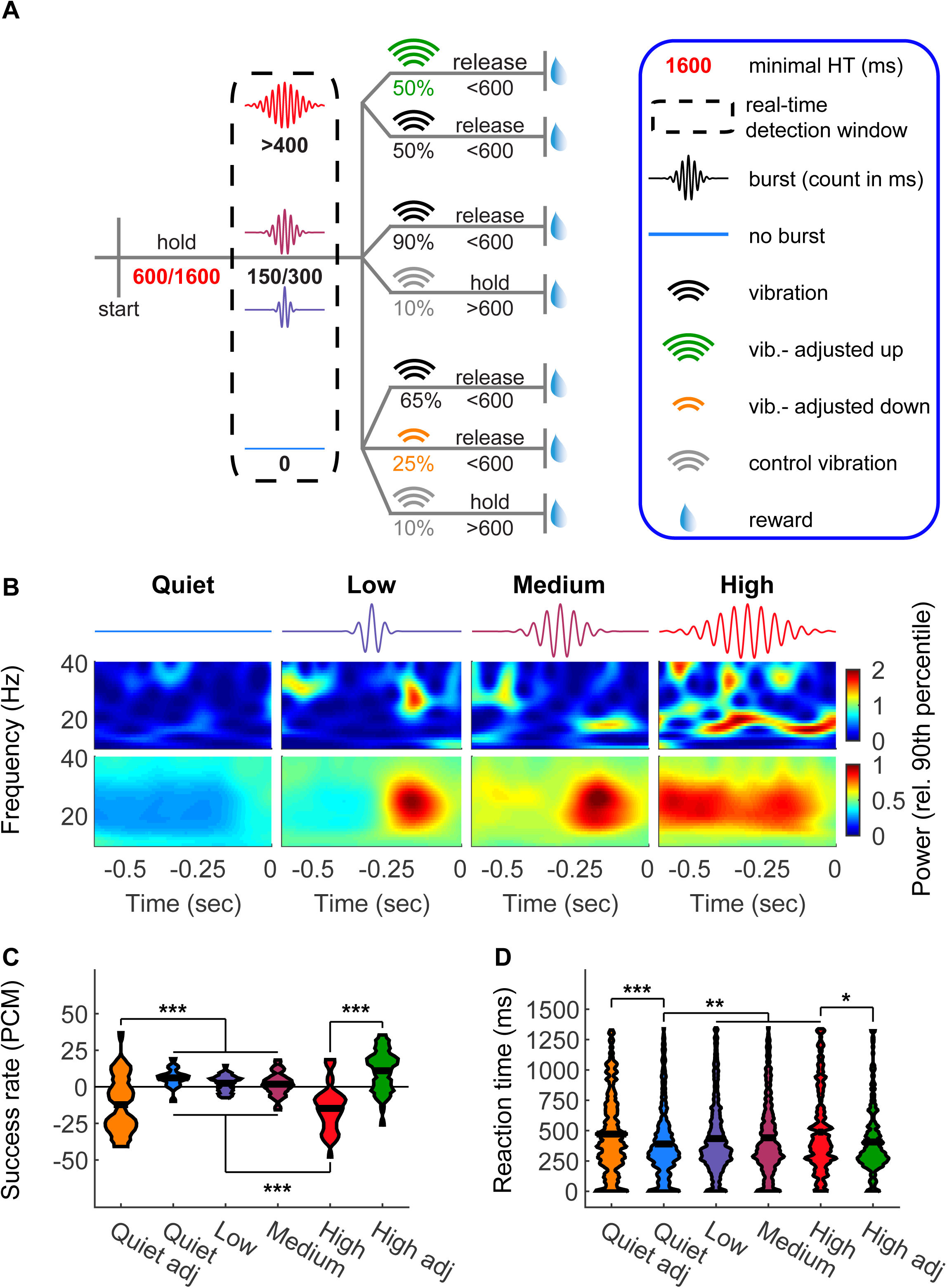
Real-time impact of beta on detection. (A) Task design. Beta-bursts were detected in real-time. A near-threshold stimulus was given upon detection of exactly 0 (quiet), 150 (low), 300 (medium), or more than 400 (high) burst samples during the detection window and the 600 ms preceding it. For adjustments, in a subset of “high” trials the stimulus amplitude was increased by 2 μm, and in a subset of “quiet” trials it was reduced by 2 μm. (B) Intrinsic brain signal triggered stimulation. Upper panels: examples of one trial per condition. Lower panels: average of all trials. Stimulus onset was at time *t*=0. (C-D) Effect of stimulus locking to intrinsic brain signal and amplitude adjustment on success rate (presented as percent change from the mean (PCM), **C**) and reaction time (**D**). The widths of the shapes in **C** indicate the distribution of sessions (N=35 from 5 rats), and in **D** the distribution of trials (N= 458, 1264, 1192, 945, 281 and 262 for quiet adjusted, quiet, low, medium, high, and high adjusted, respectively). Black horizontal lines indicate the mean. Main effects: **C**. *F*_5,204_= 23.06, *p*= 2.49*10^−48^; **D**. *F*_5,4396_= 9.29, *p*= 8.18*10^−9^. *-*p*<0.05, **-*p*<0.01, ***-*p*<0.001, ANOVA with Tukey’s critical value for multiple comparisons. See also Fig. S6.

### Bursts indicate transient synchrony of flexible cell-assemblies

Our data thus far suggest that spontaneous activity, reflected in beta-power, has an opposing role on detection to that of stimulus-evoked activity (manifested in FR and ERP, mirrored in low- and high-frequency power). But is there a direct relationship between beta oscillations and FR? Previous attempts to experimentally assess the effect of beta on stimulus-evoked FR remained controversial (Confais et al., 2020). A possible explanation for this controversy is that modulation of ongoing oscillations is not necessarily time-locked to the stimulus. This makes it difficult to identify the timing and location of the oscillations.

We found a strong positive correlation between power and burst-rate in all frequencies between 15 and 120 Hz in our task (Fig. S5). This motivated us to use the timing of the burst-power-peak as the time reference point to localize internally driven oscillations in cortical layers. We aligned bursts in each band detected on one electrode to all other electrodes, and then systematically used each electrode as the reference (Fig. S4C-E). We found that all bands had strongest ongoing power in L4. Like the evoked response (Fig. S4A), the low frequencies also showed high power in deep layers. High frequencies exhibited local synchronization as well as high power in superficial layers. In the beta band, power was maximal in L4.

The burst-peak time-referencing allowed us also to relate oscillations to spikes of single-units, pairs of units and population firing rates. Single-units demonstrated stronger LFP-spike coherence during bursts compared to epochs immediately before or after bursts in all bands (Fig. 4A, bottom). All units exhibited preference to phases of depolarization (150-210 degrees) in all bands, with phases in beta closest to the trough (Fig. 4A, top). Based on this phase preference, we defined the participating units in each burst as units firing within 30 degrees from the trough (Murthy and Fetz, 1996), and discovered that the coalition of synchronized units is dynamic. On average, each unit participated in 27.8 ± 1.12% of the beta-bursts, and each burst involved 29.2 ± 0.1% of the units recorded concurrently (mean ± 95% confidence interval). When comparing the participating units in two consecutive bursts, 45.84 ± 0.23% of the units were shared and the rest were interchanged between bursts. To investigate the effect of bursts on pair-wise synchrony, we identified significantly dependent unit-pairs based on their spike-trains (Russo and Durstewitz, 2017). We found that during bursts, the ratio of pairs with significantly dependent spike-trains from total possible pairs was significantly higher than immediately before or after bursts (Fig. 4B). An exception was high-gamma, in which the ratio of significant pairs remained high after the burst, suggesting prolongation of synchrony by the local network (Traub et al., 1999).

**Fig. 4.**
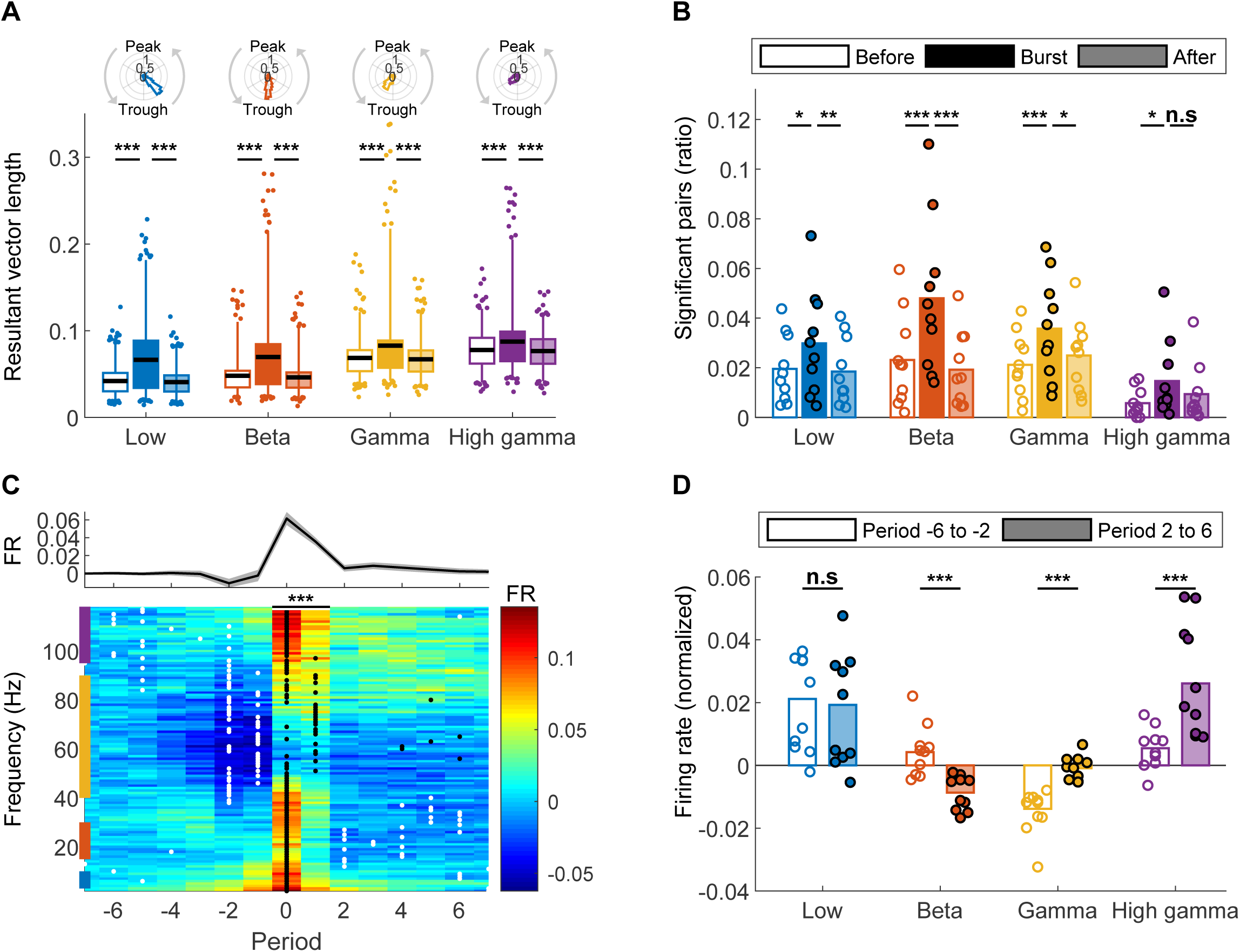
Bursts indicate transient synchrony of flexible cell-assemblies. (A) Upper panels: probability density function histograms of the mean phase at spike-times of each unit during bursts. Lower panels: the resultant vector length of spike-time-phases before (empty boxes), during (filled boxes), and after (transparent boxes) bursts. Boxes: percentiles 25 to 75. Whiskers: percentiles 2.5 to 97.5. Black line: mean. Dots: outliers. N=447 units from 5 rats. (B) The ratio of significantly correlated neuron pairs before (empty bars), during (filled bars), and after (transparent bars) bursts. Bars indicate the mean and circles individual sessions. N=10 sessions from 5 rats, on average 49.3 ± 6.1 units (1223.2 ± 257 pairs) per session. (C) Upper panel: mean ± 95% confidence interval of the normalized population firing rate (FR) per period during bursts, averaged over frequencies. Period 0 indicates the burst power maxima. Lower panel: color map of the normalized population firing rate per period and frequency. Black dots: the period with maximal FR per frequency. White dots: period with minimal FR. Colored bars on the left edge of the plot indicate the range of each band, as appears in panels A, B and D. (D) Comparison of the mean FR per band five periods before (Period −6 to −2, filled bars) and after (Period 2 to 6, empty bars) the burst peak. Bars indicate the mean and circles individual sessions. Main effects: **A**. *F*_11,4895_= 127.4, *p*= 9.38*10^−258^; **B**. *F*_2,72_= 56.5, *p*= 1.76*10^−15^;**C**. *F*_14,1624_= 47.3, *p*= 6.41*10^−110^; **D**.*F*_1,36_= 7.84, *p*= 8.15*10^−3^. * – *p*<0.05, ** – *p*<0.01, *** – *p*<0.001, n.s – not significant, repeated measures ANOVA with Tukey’s critical value for multiple comparisons.

Next, we tested the effect of bursts on FR at the population level. To this end, we extracted the population FR aligned to the timing of burst peak-power with the period of each frequency as time unit. We found that in all frequencies except low-gamma, the FR peaked at the period around the burst power maximum (period 0 in Fig. 4C). The population FR remained high for one more period before decreasing, demonstrating that FR synchrony is transient. The band-specific relationship between FR and bursts crystallizes when comparing the FR at 2-6 periods before the peak to 2-6 periods after (Fig. 4D). Low-gamma exhibited minimal FR before the burst and returned to average levels after. In the beta band, the pre-peak FR was higher and post-peak FR was lower than the mean, supporting an inhibitory effect (Schmidt et al., 2019; Seedat et al., 2020). High-gamma oscillations exhibited an increase in FR after the peak, likely due to maintenance of Pyramidal/ Interneuron Network Gamma (PING (Traub et al., 1999)) with population spikes.

Taken together, our results provide empirical evidence for transient synchrony of flexible networks during bursts, as was hypothesized in previous theoretical work (Buzsáki and Draguhn, 2004; Singer, 1994). Only in the beta-band, the transient increase in FR is followed by reduced FR, in line with the “hold” (inhibitory) function of self-generated beta (Schmidt et al., 2019; Seedat et al., 2020).

## Discussion

It is plausible that like RSN, beta-bursts represent an internally-generated co-activation of wide cortical areas. As long as this baseline activity is low, an incoming stimulus would deactivate the RSN and shift the focus from the internal state to the external environment (Deco et al., 2013). A recent study (Rumyantsev et al., 2020) demonstrated that a signal correlated over large cortical areas and orthogonal to sensory input degraded stimulus acuity, suggesting that in a high-baseline state, the internal activity can compete with the orthogonal externally evoked signal and reduce detection.

The inhibitory role of beta on behavior (Schmidt et al., 2019) and the anti-correlation between averaged beta-power and population FR (Ray et al., 2008) are consensus. However, investigation of the intrinsic relationship between FR and beta, which is important for understanding the underlying mechanism of this inhibition, led to contradicting results (Confais et al., 2020). We suggest that the reference point of the previously conducted analyses causes this controversy. Most often, the relationship between LFP and spikes is studied time-locked to the task. However, since beta is generated internally, averaging over trials time-locked to an external stimulus may blur the observed relationship. Here, by measuring the relation between spikes and LFP time-locked to the bursts, we found that in all bands, bursts indicate increased FR around the trough. Only in the beta-band bursts were followed by a reduction in firing rate, suggesting an intrinsic mechanism for the behavioral inhibition.

The difference in temporal reference point can also explain why the stimulus evoked FR does not change with performance (Lafuente and Romo, 2005) (Fig. 2b), while beta-bursts are followed by decreased FR and reduced detection. In SI, the evoked FR is time-locked to the stimulus and decodes it robustly. Beta-bursts mark a wide synchronization (Brookes et al., 2011; Hipp et al., 2012) with an inhibitory after-effect, time-locked to an internally-generated event. Since they are decoupled from the external stimulus, the FR changes are not consistently measured when looking at the stimulus time. Assuming the bursts indicate packets generated in different cortical regions and travel as a wave through cortex (Afrashteh et al., 2020; Luczak et al., 2015), it is conceivable that in higher areas, the event generated in SI is represented as a burst, and computed alongside packets generated in other sources. Future multi-site recording experiments could test this hypothesis.

In conclusion, our results reveal that beta-bursts reflect increased spontaneous activity of the RSN and predict reduced detection. This suggests that beta indicates a dynamic state that competes with detection of external stimuli. Understanding this internal state is key for the development of effective approaches of writing natural or artificial information into the brain for an optimized perception and learning as well as for neuroprosthetic approaches.

## Acknowledgments

We would like to thank Wolf Singer, Johannes Letzkus, Leor Katz, Philippe Coulon, David Eriksson and Brice delaCrompe for helpful remarks on earlier versions of the manuscript; Amiram Grinvald, James Poulet and Phillip Wisinski-Bokiniec for technical support with intrinsic-signal-imaging; and Zongpeng Sun and Irene Ayuso Jimeno for their help with preliminary experiments. This work was supported by BrainLinks-Brain-Tools, Cluster of Excellence funded by the German Research Foundation (DFG; grant no. EXC 1086), the Bernstein Award 2012 sponsored by the Federal Ministry of Education and Research (grant no. 01GQ1301), the ERC Starting Grant OptoMotorPath 338041, the Baden-Wuerttemberg Stiftung, project RatTrack as well as DFG grant nos. DI 1908/5-1, DI 1908/6-1 (all to I.D).

## Author Contributions

Conceptualization, G.K. and I.D.; Methodology, G.K.; Software, G.K.; Formal Analysis, G.K.; Investigation, G.K. and M.A.; Writing – Original Draft, G.K. and I.D.; Writing – Review & Editing, G.K. and I.D.; Visualization, G.K. and I.D.; Supervision, I.D.; Funding Acquisition, I.D.;

## Supplemental figure titles and legends

**Fig. S1. Vibrotactile detection task for freely moving rats (Related to Fig. 1).**

(A) Manipulandum. The rat had to hold the lever (I) and pull it in its preferred direction against the force of the centering magnets (II). The lever was enclosed between two walls (III) ensuring uni-manual manipulation.

(B) Complete setup. The cage was made of glass and located on a wooden surface, ensuring isolation from ground and reduced static discharge. After the holding-time, a vibrotactile stimulus was conveyed via a Piezo actuator (IV), signaling the rat to release the lever. The actuator was shielded electrically (V) to avoid electromagnetic noise in electrophysiological recordings. In a subset of trials, another shielded actuator (VI) was activated as a control. A strain gauge (VII) was used to detect the touch and to calibrate the vibration amplitude. VIII-Red cage light. IX-speaker used to deliver a clicker tone in a correct trial and white noise after errors. X-reward-delivery spout.

(C) Holding duration training progress of 8 rats. Rats had to hold the lever passed the threshold position for at least the holding time to obtain a reward. Blue lines indicate the holding duration at the beginning of each session. The holding duration was incremented automatically in small steps, until reaching the duration indicated by the red line.

(D) Holding duration during the vibro-tactile detection task. The holding time was pseudo-randomized between 600, 1600 and 2600 ms, with a 4:4:1 ratio (note the tri-modal holding-time distribution). N=6,765 trials, 40 sessions.

(E) Mean ± 95% confidence interval of the lever position (measured by the angle encoder). Time *t*=0 indicates the stimulus onset. Horizontal line at ∼1mm indicates the hold/ release threshold. Trials with holding time = 600 ms are indicated in blue, 1600 ms in purple and 2600 ms in pink.

**Fig. S2. Implant localization (Related to Figs. 1 and 2).**

(A) Intrinsic signal imaging from 8 rats. The images represent the difference in absorption intensity of 855 nm light between vibration trials and blank (no vibration) trials, relative to baseline absorption (ΔI / I) and z-normalized. Blue dashed line: assumed SI forepaw location based on published somatotopic maps^26^. Red lines: responsive area. ML: medial-lateral direction. AP: anterior-posterior.

(B) Current-sink density (CSD) map, computed as the second spatial derivative of the LFP. Time *t* = 0 indicates stimulus onset.

(C) Left, coronal section (10x) of an explanted brain after electrical lesion and NeuN staining. Area marked in red is shown on the right with higher magnification (20x). White dashed lines: estimated borders between layers. Green: lesion. Numbers: cortical layers. WM: white matter. Scale bars: left, 1000 mm, right, 200 mm.

**Fig. S3. Laminar characterization of firing rate (Related to Fig. 2).**

(A) FR of each unit (N=447 units in 10 sessions from 5 rats) during hit (left), miss (middle) and catch (right) trials. Units are sorted according to layer and timing of maximal FR. Black and white lines denote estimated limits between layers. Black and green lines denote the stimulus onset (*t* = 0) and offset (*t* = 0.3). White and blue lines: ERP phase change from de-to hyper-polarization (*t* = 0.065).

(B) Mean FR per electrode.

(C) *P*-values from statistical comparisons (two-sided *t*-test) between hit and miss (middle) and hit and control (right). Non-significant (p>0.05) time points appear in black. At stimulus onset (initial 65 ms, blue dashed line) there is a difference between hit and control, but not between hit and miss. The difference between hit and miss begins at L2/3 65 ms after stimulus onset (when the ERP changes from de-to hyper-polarization). At L4 the conditions differ after stimulus offset (0.3 sec, green dashed line).

**Fig. S4. Laminar characterization of evoked, induced and ongoing oscillations (Related to Fig. 2).**

(A) Time frequency representation (TFR) per channel of the evoked oscillations, computed by averaging the raw LFP time-locked to the stimulus, and then transferring to the frequency domain by Morlet wavelets. Numbers in magenta denote cortical layers, numbers in black the frequency in Hz. Dashed magenta lines (**A-C**) indicate estimated cortical layer borders. Time *t* = 0 indicates stimulus onset.

(B) TFR of the induced oscillations, computed by first transferring to the frequency domain by Morlet wavelets, normalizing to the median of each frequency and then averaging over trials. FOM, fraction of the median. Time *t* = 0 indicates stimulus onset.

(C) Example of localization of beta-band ongoing oscillation bursts. Time *t* = 0 indicates burst power maxima detected on electrode 5 (the reference electrode).

(D) Exemplary oscillatory bursts. Upper panel: raw LFP trace. Bursts are indicated in red. Scale bar, 0.5 mV. Lower panel: TFR of the trace on top. Bursts were defined as local maxima in the time-frequency plane, higher than the 90^th^ percentile of the peak frequency (black dots). White dashed lines: limits of the beta band.

(E) Localization of ongoing bursts in all bands and using each electrode as reference. Mean of the maximal power detected on each electrode (ordinate) during bursts in each band, as a function of the electrode used for burst detection (reference electrode, abscissa). Note that in all bands the strongest localization is at L4 (electrodes 4-6). Similar to the evoked oscillations, low frequencies were localized also in deep layers (bottom right corner, electrodes 11-15). Low and high gamma exhibit local synchronization (increased power on the diagonal). In addition, high gamma shows strong synchronization in superficial layers (electrodes 1-3). Dashed white lines: estimated borders between cortical layers.

**Fig. S5. Power dynamics are correlated with burst activity and correlated to externally-driven signal in a band-dependent manner (Related to Fig. 2).**

**(A-C)** Power dynamics and burst rate are strongly correlated for all frequencies >15 Hz. **A**. Power spectrogram, computed by transforming the LFP to the frequency domain using Morlet wavelets, normalizing to the median per frequency and averaging over hit trials. Colored lines on the left refer to band limits corresponding to color code in **C** and **D**. In all panels, time *t* = 0 indicates stimulus onset. **B**. Rate of bursts, defined similarly to the online burst rate detection algorithm, as time points with power in a certain frequency higher than the frequency above and the frequency below (in steps of 1 Hz) and higher than the 90^th^ percentile per frequency. The number of burst time points was averaged over hit trials and normalized to the median of each frequency. **C**. Power (averaged over frequencies ± 95% confidence interval indicated as shaded areas) per band with violet referring to high gamma, yellow to low gamma, orange to beta, and blue to low frequencies. Black refers to mean ± 95% confidence interval of the burst rate (BR). ρ denotes Pearson’s correlation coefficient. All data in **A-C** were recorded on electrode 5 (L4).

**(D)** LFP power correlation with externally driven signal is layer- and band-specific. Colored lines: power dynamics in L4 (electrode 5) for beta and low gamma, and in L2/3 (electrode 2) for high gamma and low frequencies. Black: Normalized population firing-rate (FR, for high gamma, low gamma and beta) or the TFR of the ERP in L2/3 (channel 3, for low frequencies).

FOM, fraction of the median, BR, burst rate, FR, population firing rate.

**Fig. S6. Trial-by-trial impact of beta-burst rate on detection (Related to Fig. 3).**

Effect of categorical segregation of beta burst rates per trial on percent change from the mean (PCM) of success rate (**A**) and reaction time (**B**). Quiet: no bursts during holding-time (HT). Low: less than a quarter of HT includes bursts. Medium: quarter to half the HT includes bursts. High: more than half the HT includes bursts. The widths of the shapes in **A** indicate distribution of sessions (N=10 from 5 rats), and in **B** the distribution of trials (N= 425, 1006, 510 and 138 for quiet, low, medium and high burst rates, respectively). Black horizontal lines indicate the mean. Main effects: **A**. *F*_3,83_= 6.68, *p*= 4.3*10^−4^; **B**. *F*_3,2075_= 10.43, *p*= 8.29*10^−7^. ** – *p*<0.01, *** – *p*<0.001, ANOVA with Tukey’s post-hoc.

## STAR Methods

### RESOURCE AVAILABILITY

#### Lead contact

Further information and requests for resources and reagents should be directed to and will be fulfilled by the Lead Contact, Ilka Diester.

#### Materials Availability

This study did not generate new unique reagents.

#### Data and Code Availability

The datasets and code used in the current study are available from the Lead Contact upon reasonable request.

### EXPERIMENTAL MODEL AND SUBJECT DETAILS

In this study we used adult (8 weeks at beginning of training) female Sprague Dawley rats (Charles-River, Sulzfeld, Germany). All procedures were in accordance with the guideline RL 2010 63 EU and were approved by the Regierungspräsidium Freiburg. We kept the rats in a reversed 12-hour light cycle, and during behavioral training gave water only during training sessions for 5 days per week and *ad libitum* over the weekend. All rats maintained >85% *ad libitum* body weight and appropriately increased weight with age.

### METHOD DETAILS

#### Behavioral setup

We developed a delayed Go/No-Go task and setup for freely moving rats (Fig. 1 and Fig. S1). For early training stages, we costume-built 4 setups for simultaneous training, controlled individually by Med-PC software (Med Associates, Fairfax, VT). Each training box included a 30×25×30 cm Plexiglas box with a grounded metal floor. A 2 × 12mm infusion cannula (1464LL, Acufirm, Dreieich, Germany) covered with a 7 mm (diameter) metal ball served as a lever. The lever was clamped to a holder 1 cm above the cage ground and between two 65 × 35 mm retractable Plexiglas walls. The holder was centered by two pairs of magnets (adhesive force 2.5 kg, model S-08-08-N, Supermagnete, Gottmadingen, Germany). We controlled the distance between magnets, and thus their force, by attaching them to M12 screws. The axis of the holder was connected to a 10-bit magnetic angle encoder (AEAT-6010, Avago Technologies, San Jose, CA), reporting the left-right position. The metal holder was connected via a high (0.5 GOhm) resistance to a 5V source, thus forming a conductive touch sensor. To deliver vibrotactile stimuli, we glued a small vibrating motor (3V ERM motor, Digikey no. 1597-1244-ND, Seeed, Shenzhen, China) to the lever. The vibrator, touch sensor, and angle encoder were controlled by an Arduino Uno (Arduino, Turin, Italy) connected via transistor-transistor-logic (TTL) to the Med Associates control cabinet. We controlled the red cage light, reward delivery infusion syringe pump (PHM-107, Med Associates), and cage speaker (ENV-224AM, Med Associates) directly with the Med Associates cabinet.

We also built a slightly modified setup for well-controlled tactile stimulation and noise free electrophysiological recordings. We used an open-top 30 × 26 ×40 cm glass cage positioned on a wood plate to ensure the rat is not grounded. We glued a 3D printed holding-ball to the lever and used a strain-gauge (SEN-14728, Sparkfun, Boulder, CO) as a touch sensor. For tactile stimulation we used a piezo actuator with 30 μm travel range (P-841.20, Physik Instrumente, Karlsruhe, Germany) powered by an E-625.S0 amplifier (Physik Instrumente). We controlled the input function to the amplifier by the electrophysiological recording system and software (RZ2 bioamp, Tucker-Davis-Technologies (TDT), Alachua, FL). To calibrate the output voltage to lever movement, we used an optical precision micrometer (optoCONTROL 2520-46, Micro-Epsilon, Uhingen, Germany). Then, we translated the lever movement to potential on the gauge-sensor and used the gauge-sensor to calibrate the lever movement before each session. Since the Piezo actuation requires high voltage (±30 V), we shielded the actuator with a light-weight 3D printed shield covered with a silver/copper shielding aerosol (#247-4251, RS Components, Morfelden-Walldorf, Germany). The shield surrounded the actuator without touching it and connected to the ground. An additional shielded piezo actuator connected to a lever served as a no-touch control. The noise and artifacts generated by the behavioral setup were <3 μV RMS (root mean square).

#### Behavioral training

We initially trained the rats to hold the lever steadily. First, we acclimated the rats to the behavioral setup for one 30 min session. Then, with the restricting walls open, we rewarded the rats with 3% sucrose water accompanied with a 12 kHz pure tone clicker upon touching and/or moving the lever. After associating touching the lever with the reward, we narrowed the gap between walls to 2 cm, and automatically increased the holding duration in steps of 10 ms after each successful hold. If a rat used its mouth or both forepaws, we manually decreased the reward size and increased the reward size upon unimanual manipulation. We inspected the preferred holding direction and paw of each rat, and hold was defined as moving and keeping the lever beyond a 1 mm threshold. Typically, the rats pulled the lever ∼5 mm, until reaching a mechanical limit (Fig. S1). The rats reached a 3 second holding duration within 10-15 sessions (Fig. S1, N=8 rats).

Next, we introduced the vibrotactile stimulus. To discourage timing, we randomized the holding duration until stimulus presentation between 600, 1600 and 2600 ms (with 4:4:1 ratio, Fig. 1 and Fig. S1), and the allowed reaction time (RT) window was automatically decreased from 2000 to 620 ms. RT shorter than 120 ms was not rewarded due to a physiological minimal detection duration (Hardung et al., 2017; Risterucci et al., 2003). To rule out a response to the vibrator sound, in 20% of the trials, we used a control vibration, outside the reach of the rats. In these trials, the rat had to continue holding for longer than the allowed RT window. The stimulus was set to 300 Hz. Throughout the behavioral task, we used the 12 kHz pure tone clicker to indicate correct trials to the rats, and white noise to indicate errors (early/late releases). After an error, at least 1 sec with the lever at center (time-out) had to pass until a new trial could be started.

#### Surgery and intrinsic signal imaging

To accurately localize the forelimb SI, while taking differences in vasculature and somatotopy between individuals into account, we utilized intrinsic-signal-imaging. This method is based on small (∼0.1%) changes in absorption of red light by active cortical areas (Grinvald et al., 2016). To detect these small changes, it is important to maintain low anesthesia with minimal pain. We used 3% isoflurane (CP-Pharma, Burgdorf, Germany) in oxygen-enriched air to anesthetize the rats and administered 10 mg kg^−1^ Carpofen (Rimadyl, Zoetis, Berlin, Germany) as analgesic. We positioned the rats in a costume-built stereotactic frame equipped with a mouth holder with palate bar (Model 929-B, David Kopf Instruments, Tujunga, CA), head-restraining mask and blunt ear bars. Before cutting the skin, we injected locally 200 μl lidocaine (0.5% w/v, bela-pharm, Vechta, Germany) over the scalp, and applied 2% lidocaine gel (Aspen Pharma, Durban, South Africa) on the affected skin after cutting. We thinned the skull above the somatosensory cortex, until brain vasculature was visible upon application of saline (0.9%). We paid special attention not to heat the brain-tissue nor cause bleeding. Then, we covered the thinned-bone with 1% agar at 37°C and covered it with a coverslip (12 mm diameter, thickness no. 1, Carl Roth, Karlsruhe, Germany). We captured a reference picture of the brain with a CMOS camera (HD1-D1312-160-CL, photon focus, Lachen, Switzerland) with tandem lenses (2x optical magnification) under power LED green illumination (530 nm, Lumileds, San Jose, CA) and controlled by an Imager 3001 system (Optical imaging, Rehovot, Israel). Then, we reduced the isoflurane concentration to 0.8-1.2%, until high-frequency, low-amplitude whisking was visible. Afterwards, we commenced the intrinsic signal acquisition under infra-red LED illumination (855 nm, Würth Elektronik, Niedernhall, Germany). We positioned a metal cannula glued to a vibrator under the forepaw of the rat, recorded the baseline signal for 4 sec at 15 frames sec^−1^ and stimulated for 4 sec. The inter stimulus interval was 17 seconds. To determine the responding area, we compared the change in absorption at frames 90-140 (stimulus) to frames 10-59 (baseline) between stimulation and control (no-vibration) trials.

Next, we increased anesthesia to 3% isoflurane and finalized a small (<1mm) craniotomy over a responsive cortical area free of big blood vessels. Without removing the dura, we inserted the 2-shanks, 32 channels laminar probes (E32+R-150-S2-L6-200-NT, ATLAS Neuroengineering, Leuven, Belgium) at a speed of 5 μm per second using a costume-made motorized inserter to a final depth of 2300 μm. While holding the probe with a vacuum holder (ATLAS), we applied a sealant (Kwik-Cast, World Precision Instruments, Sarasota, FL) over the craniotomy and thinned skull and UV-cured cemented the probe to the skull (RelyX, 3M, Saint Paul, MN). For increased stability and reduced noise, we also cemented the costume-made electrode interface board and protected it with a costume-made implantable metal M18 screw and cup (Froelich AG, Untersiggenthal, Switzerland). Rats were given >7 days of recovery before continuation of behavioral and electrophysiological recordings.

#### Data acquisition

We sampled the broadband signal at 25 kHz using a digital head stage (ZD32, TDT). Spiking activity was bandpass filtered between 300 and 5,000 Hz, and LFP was low pass filtered at 500 Hz and down sampled to 1 kHz. The Med Associates system registered behavior at 100 Hz and synchronized with the electrophysiological signal via TTL communication. For real-time LFP burst detection (Fig. 3) we used a recently reported method (Karvat et al., 2020). Briefly, we filtered the raw signal by narrowband bandpass filters centered on frequencies 1 to 32 Hz in steps of 1 Hz. We estimated power as the square of the amplitude at turning points of the filtered LFP. Bursts were defined as power in the frequencies 15-30 Hz which exceeded the 90^th^ percentile (calculated online over the passing 2 min) and was higher than the frequency above and frequency below. Detected bursts were sent to the behavioral controller, which summed them during trials. After the minimal holding duration, we allowed 1 sec window of beta-burst detection. Time points with bursts were summed over the 600 ms before and during the detection window, and the vibrotactile stimulus was sent upon detection of exactly 0 (quiet), 150 (low), 300 (medium), or more than 400 (high) burst samples. Half of the high burst-rate trials and quarter of quiet trials were pseudo-randomly chosen for adjusted cue amplitude. To investigate the effect of the minimal detectable changes in amplitude in our system (2μm), we reduced the vibration amplitude to the minimal level resulting in d’ of 1 at quiet trials for each rat (8.8 ± 1.36 μm, d’=0.99 ± 0.18, mean ± 95% confidence interval, n=5 rats).

#### Data analysis

To calculate the event-related-potentials (ERP, Fig. 2A), we aligned the raw LFP to the stimulus onset time and averaged over trials (n=10 sessions, 5 rats). To measure the current-source-density (CSD, Fig. S2B) we first calculated the second spatial derivative of the LFP and then averaged over trials. For firing-rate analyses, we sorted the broadband signal into units using KiloSort (Pachitariu et al., 2016). Then we inspected each cluster and defined units based on wave shape and existence of refractory periods. We estimated the firing rate (FR) of each unit by a 25 ms boxcar filter and the population FR as the population-mean of the *z*-normalized FR of each unit at each time point. We defined units as modulated if they exhibited significantly higher FR at the period [0 500] ms (stimulus present at [0 300]) compared to a baseline of [-1000 0), and also a significant difference between stimulus and control trials.

We transformed LFP data from the time domain to the frequency domain by convolving it with Morlet wavelets seven cycles wide in each frequency using the Fieldtrip toolbox (Oostenveld et al., 2011) for Matlab (Mathworks, Natick, MA, version R2018a). To assess the evoked response (Fig. S4A) we used the ERP as input signal. For induced response calculations (Fig. 2C-D, Figs. S4B and S5) we transformed the raw LFP (of the whole session) to the frequency domain, then removed artefactual segments defined as timestamps in which the LFP exceeded 1mV and 500 ms before and after them (Karvat et al., 2020), and normalized each frequency to the 90^th^ percentile of its artifact-free power. For comparison between behavioral outcomes (Fig. 2d) and correlation with FR or ERP (Fig. S5D), we chose the most modulated channel for each band, defined as the channel with the biggest power range during the period [-300,300] ms (stimulus present at [0 300]).

For comparison with power dynamics, we computed offline burst-rate in a similar manner to the online algorithm (Fig. S6), from one electrode located in L4. We defined a time point as having a burst in a specific frequency if the power in that frequency was higher than the power in the frequency above and the frequency below (in steps of 1 Hz). In addition, the power had to exceed the 90^th^ percentile of that frequency (calculated over the whole session). The number of time points having bursts was averaged over hit trials and normalized to the median rate of each frequency. To calculate the laminar localization of ongoing oscillations (Fig. S4C-E), we defined bursts as peaks in the time-frequency plane (Shin et al., 2017) exceeding the 90^th^ power percentile (Fig. S4D). We detected bursts based on the LFP-power in one electrode (the reference electrode), and averaged the power at the interval −200 to 200 ms relative to the peak on all electrodes. We repeated the analysis taking each electrode as reference and for each band (low: 3-10 Hz, beta: 15-30 Hz, low-gamma: 45-90 Hz and high-gamma: 95-120 Hz).

To investigate the effect of bursts on synchrony (Fig. 4A-B), we defined the burst-duration as the time interval *t*_[0,T]_ around a peak in which the power exceeded the 90^th^ percentile of the frequency at peak. Then, for each burst, we defined the time interval *t*_[-T-1,-1]_ as a “before burst” interval, and the time interval *t*_[T+1,2T+1]_ as an “after burst” interval. We combined these intervals per band into time vectors to mask all spikes per unit, and extracted the resulting vector length of the LFP phase at spike times. For phase histograms (Fig. 4A, top) we used the circular mean of the phase of each unit. To investigate pairwise synchrony (Fig 4B), we adopted a parametric statistic approach to test for pairwise dependencies of spike trains with correction for non-stationarity (Russo and Durstewitz, 2017). We input the spike-trains masked for before, during and after bursts into the algorithm provided by Russo and Durstewitz (Russo and Durstewitz, 2017), and defined the ratio of pairs per session as the number of significantly (*p* < 0.05) synchronized pairs divided by the number of all possible pairs in the sessions.

For investigation of the period-by-period effect of bursts on population firing rate (Fig. 4C-D), we defined the peak time ± half a period per frequency as Period 0, and extracted the population FR 10 periods before and 10 periods after each burst-peak. We then averaged the FR in each period, and normalized to a baseline of period – 10 to −8. For the before-after burst comparison (Fig. 4D), and since the period immediately after the peak showed increased FR, we used the periods −6 to −2 as “before” and periods 2 to 6 as “after”.

We presented success rates of different burstiness categories (Fig. 3C and Fig. S6A) as percent change from the mean (PCM). As reaction-times (Fig. 3D and Fig. S6B) we took the duration between vibration initiation and the lever passing the hold-threshold.

#### Histology

To validate the depth of the electrode used for real-time burst detection, in the day before sacrificing the rats we created an electrical lesion by applying a 10 μA current for 14 seconds (Karlsson et al., 2012) through this electrode using a constant current stimulator (STMISOLA, Biopac, Goleta, CA). We sacrificed the animals by administration of 800 mg/kg sodium pentobarbital (Narcoren, WDT, Garbsen, Germany) and perfused transcardially with Phosphate-Buffered Saline (PBS) and ice-cold 4% paraformaldehyde. Two days after extracting the brains, we transferred them into a solution of 30% w/v sucrose in water at 4°C for equilibration. Then, we sectioned the brains into 50 μm thin slices on a freezing cryostate (SM 2010R, Leica, Wetzlar, Germany). To delineate laminar boundaries, we blocked the slices with 5% Bovine Serum Albumin (BSA) in PBS for 1 hour and incubated in anti-NeuN primary antibody (rabbit polyclonal, Millipore catalog #ABN78, Merck-Millipore, Darmstadt, Germany, diluted 1:500 in PBS). Afterwards, we incubated the slices for 3 hours with the secondary antibody (CY3 rabbit anti-mouse, AP160C, diluted 1:250 in PBS) washed in PBS for 3×10 minutes. We obtained images using an Axioplan 2 microscope (Zeiss, Oberkochen, Germany), using Epiplan Neofluar 10x HD DIC (NA 0.30, air) and Plan Apochromat 20x M27 (NA 0.80, air) objectives.

### QUANTIFICATION AND STATISTICAL ANALYSIS

We tested the difference between Hit and False-alarm rates (Fig. 1E) with two-sided Wilcoxon rank sum test. To measure the difference between trial types (hit, miss and control) in FR and LFP over time (Fig 2A, B, and 2D), we divided the analyzed period into 25 ms time bins, and performed a repeated-measures ANOVA with Tukey’s critical value for multiple comparisons. Similarly, we used ANOVA for repeated measures to test the effect of bursts on synchrony (before, during and after burst, Fig. 4A, B) and the effect of periods relative to peak on FR (Fig. 4C, D). To assess the effect of burst rate on success rate (Fig. 3C and Fig. S6A) and reaction-time (Fig. 3D and Fig. S6B) we used a one-way analysis of variation (ANOVA) with Tukey’s adjustment for multiple post-hoc comparisons. All statistical tests were performed in Matlab, with significant level of α = 0.05 and statistical details can be found in Results and figure legends.

